# MR1-restricted MAIT cells from the human lung mucosal surface have distinct phenotypic, functional, and transcriptomic features that are preserved in HIV infection

**DOI:** 10.1101/2020.11.19.389858

**Authors:** Sharon Khuzwayo, Maphe Mthembu, Erin W. Meermeier, Sanjay M. Prakadan, Samuel W. Kazer, Thierry Bassett, Kennedy Nyamande, Dilshaad Fakey Khan, Priya Maharaj, Mohammed Mitha, Moosa Suleman, Zoey Mhlane, Dirhona Ramjit, Farina Karim, Alex K. Shalek, David M. Lewinsohn, Thumbi Ndung’u, Emily B. Wong

**Affiliations:** Africa Health Research Institute, Durban, South Africa; School of Laboratory Medicine and Medical Sciences, University of KwaZulu-Natal, Durban, South Africa; Department of Pulmonary and Critical Care Medicine, Oregon Health and Science University, Portland, Oregon, United States of America; The Ragon Institute of MGH, MIT, and Harvard University, Cambridge, Massachusetts, United States of America; Institute for Medical Engineering & Science (IMES), Department of Chemistry, and Koch Institute for Integrative Cancer Research, MIT, Cambridge, Massachusetts, United States of America; Broad Institute of MIT and Harvard, Cambridge, Massachusetts, United States of America; Department of Pulmonology, Inkosi Albert Luthuli Hospital, Durban, South Africa; Department of Pulmonology and Critical Care, Nelson R. Mandela School of Medicine, University of KwaZulu-Natal, Durban, South Africa; Department of Molecular Microbiology and Immunology, Oregon Health and Science University, Portland, Oregon, United States of America; Department of Research, VA Portland Health Care Center, Portland, Oregon, United States of America; HIV Pathogenesis Programme, The Doris Duke Medical Research Institute, University of KwaZulu-Natal, Durban, South Africa; Max Planck Institute for Infection Biology, Berlin, Germany; Division of Infection and Immunity, University College London, London, United Kingdom; Division of Infectious Diseases, Massachusetts General Hospital, Boston, Massachusetts, United States of America; Department of Medicine, University of Alabama at Birmingham, Alabama, United States of America

## Abstract

Mucosal associated invariant T (MAIT) cells are a class of innate-like T cells that utilize a semi-invariant αβ T cell receptor to recognize small molecule ligands produced by bacteria and fungi. Despite growing evidence that immune cells at mucosal surfaces are often phenotypically and functionally distinct from those in the peripheral circulation, knowledge about the characteristics of MAIT cells at the lung mucosal surface, the site of exposure to respiratory pathogens, is limited. HIV infection has been shown to have a profound effect on the number and function of MAIT cells in the peripheral blood, but its effect on lung mucosal MAIT cells is unknown. We examined the phenotypic, functional, and transcriptomic features of MR1 restricted MAIT cells from the peripheral blood and bronchoalveolar compartments of otherwise healthy individuals with latent *Mycobacterium tuberculosis (Mtb)* infection who were either HIV uninfected or HIV infected. Peripheral blood MAIT cells consistently co-expressed typical MAIT cell surface markers CD161 and CD26 in healthy individuals, while paired bronchoalveolar MAIT cells displayed heterogenous expression of these markers. Bronchoalveolar MAIT cells produced lower levels of pro-inflammatory cytokine IFN-γ and expressed higher levels of co-inhibitory markers PD-1 and TIM-3 than peripheral MAIT cells. HIV infection resulted in decreased frequencies and pro-inflammatory function of peripheral blood MAIT cells, while in the bronchoalveolar compartment MAIT cell frequency was decreased but phenotype and function were not significantly altered. Single-cell transcriptomic analysis demonstrated greater heterogeneity among bronchoalveolar compared to peripheral blood MAIT cells and suggested a distinct subset in the bronchoalveolar compartment. The transcriptional features of this bronchoalveolar subset were associated with atypical MAIT cells and tissue repair functions. In summary, we found previously undescribed phenotypic and transcriptional heterogeneity of bronchoalveolar MAIT cells in healthy people. In HIV infection, we found numeric depletion of MAIT cells in both anatomical compartments but preservation of the novel phenotypic and transcriptional features of bronchoalveolar MAIT cells.

## Introduction

Mucosal associated invariant T (MAIT) cells are a relatively recently described group of innate-like T cells that recognise antigen presented by the highly conserved MHC class I-related (MR1) protein [1, 2]. MAIT cells classically recognize and respond to a variety of bacteria, through MR1-restricted recognition of bacterially derived riboflavin metabolites [3–5]. MAIT cells also recognize and respond to *Mycobacterium tuberculosis (Mtb)* although it is unclear whether this interaction is mediated via the recognition of intermediates generated by the riboflavin biosynthesis pathway [6]. MAIT cells have been found to be enriched and display pro-inflammatory function in the lungs of people with active tuberculosis (TB) [7]. In mouse models, MAIT cells in the lungs have been shown to play a role in the establishment of a coordinated response to respiratory pathogens [8]. It has been demonstrated that MAIT cells make up a large proportion of *Mtb*-reactive CD8+ T cells in the peripheral blood of humans, suggesting their potential importance in anti-TB immunity [3, 9]. Nonetheless, the specific role of MAIT cells in protection against human respiratory infections, including TB, remains unclear [10–12]. MAIT cells are also able to respond to viral infection, the recognition of which is mediated by IL-18 and IL-12 stimulation [13, 14] rather than by ligand-driven T cell receptor mediated response. HIV infection is known to alter immunity to respiratory bacterial and fungal pathogens including Mtb, the causative pathogen of TB, which is the leading cause of death for people living with HIV worldwide [15]. In HIV infection, MAIT cells are depleted in peripheral blood and are functionally exhausted [16, 17]. The impact of HIV on MAIT cells at the lung’s mucosal surface is incompletely understood [18].

In this study, we aimed to understand the unique features of MAIT cells in the bronchoalveolar compartment of healthy HIV-negative humans with latent *Mtb* infection by using surface phenotyping, functional analysis, and transcriptomics. We then compared MAIT cells from HIV-negative individuals to those from HIV-positive individuals to better understand the impact of HIV infection on MAIT cells at the respiratory mucosal surface.

## Methods

### Study participants

Bronchoalveolar lavage (BAL) fluid and matching peripheral blood samples were collected from participants undergoing research (n=37) or clinically indicated (n=17) bronchoscopies at Inkosi Albert Luthuli Central Hospital (IALCH) in Durban, South Africa. All participants completed written informed consent for study procedures. Ethical review and approval of the study protocol was received from the University of KwaZulu-Natal (UKZN) Biomedical Research Ethics Committee (BREC) (protocol numbers BF503/15 and BE037/12) and the Partners Institutional Review Board. In the research bronchoscopy cohort, there were two groups of participants: 1) “Healthy controls” defined as HIV-negative (negative HIV ELISA and undetectable HIV viral load) and with latent TB infection (LTBI) as indicated by a positive QuantiFERON-TB Gold Plus (QFT-Plus) assay (Qiagen) (IFN-γ > 0.35 IU/mL after subtracting background), and 2) “HIV-positive participants” who were newly diagnosed with a positive HIV ELISA and detectable HIV viral load, antiretroviral therapy naive, with good functional status, free from symptoms of respiratory infection (cough, fever, shortness of breath) and with confirmed LTBI by the QFT-Plus assay. Participants with a history of TB as well as those with signs that could be considered consistent with active TB by chest X-ray were excluded from the research bronchoscopy cohort. Participants undergoing clinically indicated bronchoscopies had their HIV status defined by HIV ELISA and viral load, and had respiratory infections excluded by negative bronchoalveolar bacterial, fungal and mycobacterial cultures at the time of enrolment. The characteristics of the study participants from both cohorts are shown in **Table 1**.

**Table 1:** Demographic and clinical characteristics of participants from clinically-indicated (clinical) and research bronchoscopy study cohorts.

|  | Research bronchoscopy cohort |  |  | Clinical bronchoscopy cohort |  |  |
| --- | --- | --- | --- | --- | --- | --- |
|  | HIV- | HIV+ | *P*-value | HIV- | HIV+ | *P*-value |
| **N (%)** | 23 (62.2%) | 14 (37.8%) | 0.3333 | 8 (47%) | 9 (53%) | 0.6667 |
| **Age, years (IQR\*)** | 32 (25 - 36) | 30 (21.75 - 35.25) | 0.4998 | 45 (29.25 - 54.75) | 34 (32 - 51) | 0.833 |
| **Sex, number of females (%)** | 16 (69.6%) | 9 (64.3%) | 0.3333 | 2 (25%) | 4 (44.4%) | 0.3333 |
| **Median viral load, copies/mL (IQR\*)** | Undetectable | 25 229 (9768 - 60 331) | N/A | Undetectable | 304 (<20 - 480 000) | N/A |
| **CD4 count, cells/μL (IQR\*)** | 1005 (833.3 - 1308) | 445 (193.8 - 615.3) | <0.0001 | ND | 276 (96 - 589) | N/A |

\*Interquartile range.
ND - not done

### Lymphocyte preparation

Bronchoscopy was performed by pulmonologists at Inkosi Albert Luthuli Hospital in Durban, South Africa according to standard protocols and in accordance with local standard of care [7]. Directly following the procedure, bronchoalveolar lavage (BAL) fluid was either combined 1:1 with complete RPMI (cRPMI) media (supplemented with 10% fetal bovine serum, 200 mM L-glutamine, 10 000 U/mL penicillin-streptomycin and 250 μg/mL amphotericin) or transported undiluted back to the laboratory on ice. All samples were processed within 3 hours of collection. BAL fluid was filtered through a 40 μm strainer (BD Pharmigen), centrifuged, and the pelleted lymphocytes resuspended in cRPMI media for immediate assay or cryopreserved. In parallel, peripheral blood mononuclear cells (PBMCs) were isolated from whole blood using the Histopaque^®^ gradient centrifugation method, according to the manufacturer’s instructions (Sigma-Aldrich) and suspended in cRPMI for immediate use or cryopreserved.

### Flow cytometry and sorting

Surface staining was performed on paired BAL lymphocytes and PBMCs using 1:500 of either MR1 5-OP-RU (PE) or MR1 6-FP (PE) tetramer (NIH Tetramer Core Facility), 1:800 of a fixable viability dye (Live/Dead Aqua, Invitrogen) and one of two surface staining antibody master mixes. The first included 1:100 of anti-CD3 PE-CF594 (UCHT1, BD Biosciences), 1:50 of anti-CD4 BV711 (OKT4, BioLegend), 1:200 of anti-CD8 APC-H7 (SK1, BD Biosciences), 1:50 of anti-CD26 FITC (BA5b, BioLegend), 1:50 of anti-CD161 PE-Cy7 (HP-3G10, BioLegend), and 1:50 of anti-PD-1 BV421 (EH12.1, BD Biosciences). The second antibody master mix used included 1:50 of anti-CD3 BV650 (OKT3, BioLegend), 1:50 of anti-CD4 BV711 (OKT4, BioLegend), 3:100 of anti-CD8 PE-Texas red (3B5, Thermo Fisher), 1:50 of anti-CD26 FITC (BA5b, BioLegend), 1:100 of anti-CD161 PE-Cy7 (HP-3G10, BioLegend), 1:50 of anti-PD-1 BV421 (EH12.1, BD Biosciences) and 1:50 of anti-TIM-3 BV785 (F38-2E2, BioLegend). Tetramer staining was performed in the dark at room temperature for 40 minutes, followed by 20 minutes of staining with the fixable viability dye and surface antibodies at 4°C. All samples were acquired on the FACSAria™ III (BD Biosciences). Rainbow Fluorescent Particles (BD Biosciences) and applications settings in BD FACSDiva v7.0 were used to correct for day-to-day variations in instrument performance. The data were analysed using FlowJo™ (v10.4). MR1 5-OP-RU tetramer-positive cells were gated on from the population of live (Aqua Live/Dead-negative), single lymphocytes (based on forward and side scatter) that were CD3+CD4-(except where otherwise indicated). The MR1 6-FP tetramer was used to define the positive gate for the MR1 5-OP-RU tetramer, and gates for PD-1 and TIM-3 staining determined using fluorescence minus one controls.

### Intracellular cytokine staining assay

When sufficient cells were available, freshly isolated matched BAL lymphocytes and PBMCs were simultaneously stimulated using PMA (25 ng/mL) and ionomycin (500 ng/mL) for 6 hours at 37°C in 96-well microplates. After 6 hours, plates were washed in phosphate buffered saline (PBS) before cells were stained with 1:500 of either MR1 6-FP (PE) or MR1 5-OP-RU (PE) tetramer (NIH Tetramer Core Facility). Tetramer staining was performed in the dark at room temperature for 40 minutes, followed by surface staining at 4°C for 20 minutes. The surface staining antibody master-mix included: 1:25 of anti-CD4 BV711 (OKT4, BioLegend), 1:100 of anti-CD8 APC-H7 (SK1, BD Biosciences), 1:50 of anti-CD26 FITC (BA5b, BioLegend), 1:50 of anti-CD161 PE-Cy7 (HP-3G10, BioLegend) and 1:800 of a fixable viability dye (Live/Dead Aqua, Invitrogen). Cells were then fixed (Fix & Perm Medium A, Invitrogen) at room temperature in the dark for 15 minutes and permeabilised (Fix & Perm Medium B, Invitrogen) for 20 minutes at 4°C. Intracellular antibodies were added during permeabilization and included 1:50 each of: anti-CD3 AlexaFluor700 (UCHT1, BioLegend), anti-IFN-γ PE-Dazzle594 (4S.B3, BioLegend), anti-granzyme B AlexaFluor647 (GB11, Biolegend) and anti-IL-17 BV421 (BL168, BioLegend). Cells were PBS washed, acquired using the LSRFortessa™ (BD Biosciences) and the data then analysed using FlowJo (v10.4).

### RNA isolation and sequencing

MR1 tetramer-positive cells (gated as stated above) from paired BAL fluid and PBMCs were either bulk sorted in 100-cell mini-populations or single-cell sorted using the FACSAria™ III (BD Biosciences) in purity mode into 96-well microplates containing 10 μL of 1% 2-mercaptoethanol RLT buffer (Qiagen) and stored at −80°C. RNA capture and library preparation for single cells was performed using the Smart-Seq2 approach as described by Trombetta *et al.* [19]. Briefly, MR1 tetramer-positive cells were lysed, RNA isolated using magnetic SPRI bead isolation (Agencourt RNAClean XP, Beckman Coulter) and cDNA then synthesized from full-length RNA sequences. Whole transcriptome amplification (WTA) was then performed, and the WTA product quality determined by assessing fragment size using the BioAnalyzer (Agilent Technologies) and the concentration quantified using the Qubit® fluorometer and assay kit (Thermo Fisher). Nextera XT libraries were then constructed, pooled and purified using DNA SPRI beads (Agencourt AMPure XP, Beckman Coulter) before sequencing was performed on the NextSeq500 (Illumina).

### MAIT cell cloning

MAIT cell cloning was performed as described by Cansler *et al.* [20]. Briefly, MR1 tetramer-positive T cells were sorted from cryopreserved bronchoalveolar lavage cells and rested overnight. Limiting dilution assay was performed to seed single cells in 96-well plates in RPMI media containing 10% human serum (HuS), 2% L-glutamine, 0.1% gentamycin and irradiated feeder cells (lymphoblastoid cell lines and peripheral blood mononuclear cells). Media was supplemented with recombinant human (rh)IL-2 (5 ng/mL), rhIL-7 (0.5 ng/mL), rhIL-12 (0.5 ng/mL), rhIL-15 (0.5 ng/mL) (BioLegend) and anti-CD3 (0.03 μg/mL) (eBiosciences). Plates were assessed weekly for growth with clones taking 2 – 3 weeks to grow. Clones were taken forward for subsequent analyses only from plates which had growth in approximately less than 30% of the wells. Resulting clones were subjected to surface MR1 5-OP-RU tetramer and monoclonal antibody staining with 1:200 of Live/dead viability dye (Aqua, Life Technologies), 1:25 of anti-CD3 PerCP/Cy5.5 (UCHT1, BioLegend), 1:25 of anti-CD4 BV785 (OKT4, BioLegend), 1:50 of anti-CD8 (FITC, RPA-T8) and 1:25 of anti-CD26 PE-Cy7 (BA5b, BioLegend), as well as IFN-γ ELISPOT assay to confirm MAIT cell identify and clonal purity.

### IFN-γ ELISPOT assay

ELISpot plates were coated with 10 μg/mL anti-human IFN-γ coating antibody (Mabtech, clone 1-D1K) and incubated overnight at 4C°. After washing with 1x PBS, plates were blocked for 1 hour at room temperature using 10% HuS RPMI before wild type (WT) A549 and MR1^−/−^ A549 lung epithelial cells [21] were added. *Mycobacterium smegmatis (M.smeg)* was added at an MOI of 1:3 and the infection allowed to proceed for 2 – 3 hours before the addition of the suspected MAIT cell clones. A PHA control (10 μg/mL) was included and the plate incubated overnight at 37C°. The following morning plates were washed with 1x PBS-T before being coated with 3 μg/mL streptavidin-ALP secondary antibody (Mabtech, clone 7-B6-1) and incubated for 2 hours at room temperature. Detection substrate was added after performing 1x PBS-T washes and spots allowed to develop. Final deionized (DI) water rinses were performed, the plates dried for 45 minutes and the spots then quantified using an ELISpot reader (Autoimmun Diagnostika GmbH) with accompanying software.

### DNA extraction and T cell receptor sequencing

DNA was extracted from MAIT cell clones using the Qiagen DNeasy mini-kit (according to manufacturer’s instructions) and the DNA then quantified using the ND-1000 (NanoDrop®, Thermo Scientific). Paired TCRα and TCRβ sequencing was performed using the ImmunoSEQ assay (Adaptive Biotechnologies). CDR3α sequences were then assessed using an online tool, MAIT Match (http://www.cbs.dtu.dk/services/MAIT_Match), which compares sequences to reference MAIT cell sequences in the database and calculates similarity scores from 0 (mismatch) to 1 (perfect match) [7].

### Statistical and bioinformatic analyses

All statistical analyses were performed on Prism 8.0.0 (GraphPad Software) using Mann-Whitney *U* tests to assess differences between compartments (peripheral blood vs BAL fluid) or between disease state (HIV-negative vs HIV-positive). Data are represented as medians and interquartile ranges. Single-cell and bulk RNA sequence data was demultiplexed with bcl2fastq v2.17, aligned to the human genome using TopHat and the counts were estimated using RSEM [22]. Differential gene expression analysis and data visualisation of the single-cell data was performed using Seurat R package (3.0.1) [23], while DESeq2 R package (1.16.1) [24] and Prism 8.0.0 were used for the bulk RNA-sequencing analysis. Genes were considered differentially expressed when fold change > 1 or < −1 and FDR q-value < 0.05 (Benjamini-Hochberg correction).

## Results

### Lung mucosal MAIT cells are phenotypically heterogenous

We first assessed the frequency of MAIT cells, which we defined as CD3+CD4-MR1 5-OP-RU tetramer-positive cells (**Figure 1A**), in the two anatomical compartments, and found that in the healthy HIV-negative participants without any active respiratory infection MAIT cells frequencies in peripheral blood were similar to those in the bronchoalveolar compartment (*P* = 0.3027) (**Figure 1B**). Paired surface staining of MAIT cells from the peripheral blood and bronchoalveolar compartment of healthy controls using the surface markers CD161, a C-type lectin, and the dipeptidyl peptidase CD26, which are typically highly expressed by MAIT cells [25, 26], demonstrated that nearly all the peripheral blood MR1 tetramer-positive cells expressed high levels of these markers, but showed unexpected heterogeneity in expression of both in the bronchoalveolar compartment (characteristic example shown in **Figure 1C**). Assessment of all the healthy HIV-negative participants from the research bronchoscopy cohort found that the vast majority of peripheral blood MAIT cells had the CD161++CD26++ phenotype [median of 94.2% and interquartile range (IQR) 87.9 – 97.9%] and that this phenotype was significantly less frequent (64.95%, 39.0 – 77.88%) among MR1 tetramer-positive cells derived from the bronchoalveolar fluid (*P* = 0.0002) (**Figure 1D**). Although CD161-negative and CD26-negative staining MR1 tetramer-positive cells were very rare in the peripheral blood, MR1 tetramer-positive cells in the bronchoalveolar compartment contained cells falling into these categories at detectable frequencies with medians of 13.5% for CD161-CD26++ (IQR: 7.41 – 27.23%), 10.25% for CD161-CD26-(IQR: 1.86 – 18.58%) and 7.66% for CD161++CD26-(IQR: 1.94 – 15.7%) cells. These phenotypic subpopulations were significantly more frequent in the bronchoalveolar compartment compared to matched peripheral blood samples (*P* = 0.0034, 0.0084 and 0.0013, respectively).

**Figure 1:**
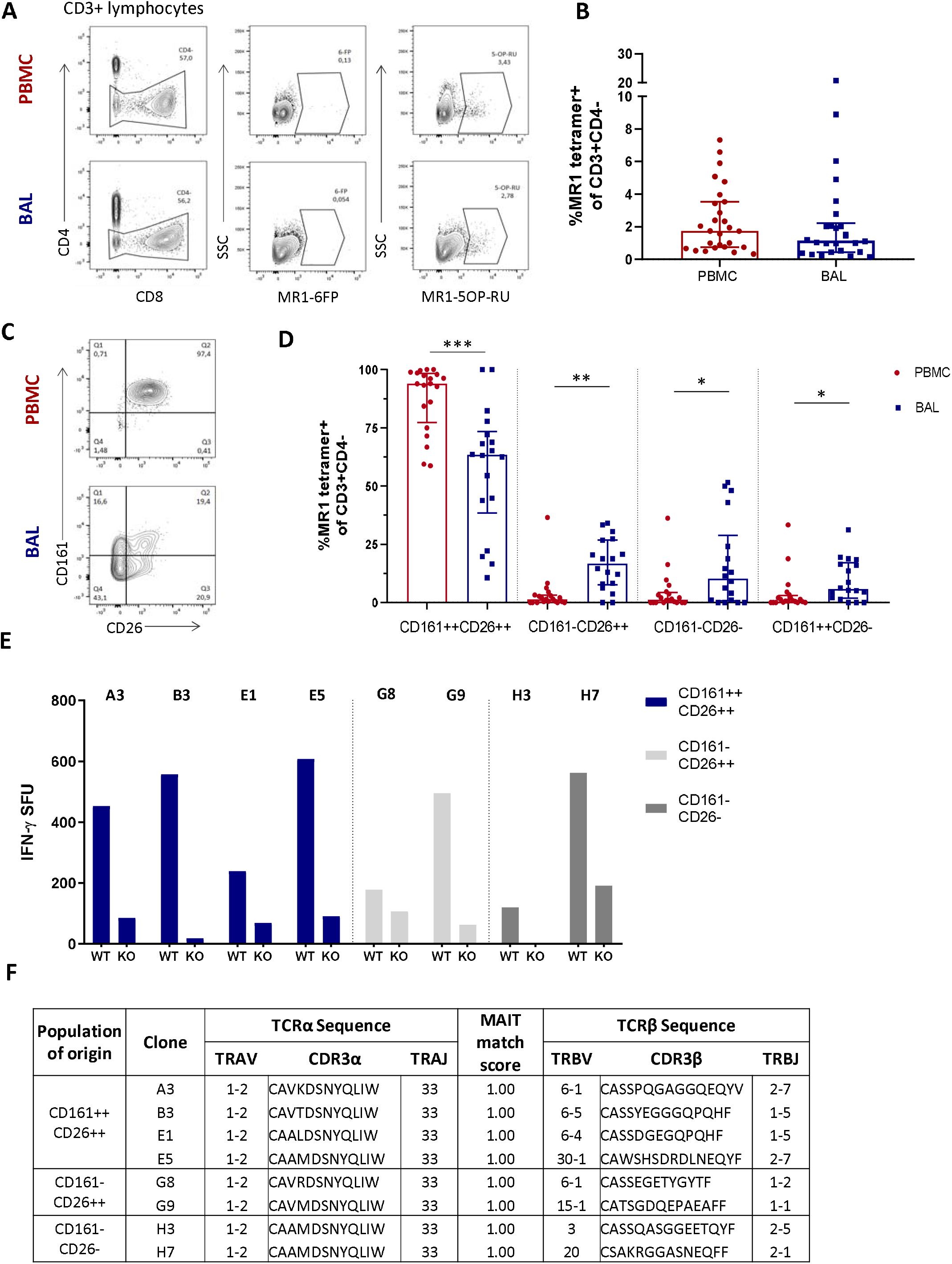
Phenotypic heterogeneity of bronchoalveolar MAIT cells in healthy individuals. (A) Gating strategy used to define CD3+CD4-MR1 5-OP-RU tetramer-positive cells (using the MR1 6-FP tetramer as a control) and (B) MAIT cell frequency in the peripheral blood (PBMC) and bronchoalveolar (BAL) compartments. (C) Representative CD161 and CD26 staining of peripheral blood and bronchoalveolar CD3+CD4-MR1 tetramer-positive MAIT cells and (D) frequency of MAIT cell phenotypic subpopulations in peripheral blood (red) and bronchoalveolar lavage (BAL) fluid (blue) (*P* = 0.0002, 0.0034, 0.0084 and 0.0013 respectively). MR1 tetramer-positive T cells with atypical MAIT cell phenotype were cloned from the bronchoalveolar compartment and IFN-γ ELISpots performed to confirm MAIT cell identity. (E) MR1-dependent function of the resulting 8 bronchoalveolar MAIT cell clones (WT = M. smeg infected wild type A549 cells; KO = M. smeg infected MR1^−/−^ A549 cells). (F) T cell receptor sequencing of bronchoalveolar MR1 tetramer+ T cell clones grouped by phenotypic subpopulation of interest. All clones possessed the MAIT cell semi-invariant TRAV1-2/TRAJ33 alpha-chain and had MAIT match scores of 1.00 based on CDR3α sequences. Bars represent medians and error bars represent interquartile ranges. Statistical difference was determined using the Mann-Whitney *U* test.

There have been reports of CD161-low/negative MAIT cells [17, 27], but because MAIT cells with a CD161-negative and CD26-negative phenotype have not been previously described, we sought to confirm the identify of these phenotypically heterogenous MR1 tetramer-positive cells by cultivating T cell clones from these phenotypic subpopulations to allow assessment of their functional characteristics and T cell receptor usage. We therefore selected a participant with prominent CD161 and CD26 heterogeneity in the bronchoalveolar compartment and cloned cells from the CD161++CD26++, CD161-CD26++ and CD161-CD26-subpopulations of bronchoalveolar T cells. Of the eight clones successfully cultivated, four were from the CD161++CD26++ subpopulation, two were from the CD161-CD26++ subpopulation and two were from the CD161-CD26-subpopulation. Functional assessment showed that all the cultivated clones, regardless of the CD161/CD26 subpopulation from which they had been sorted, had MR1-restricted IFN-γ production in response to *M.smeg* infection of non-HLA-matched antigen presenting cells, consistent with MAIT cell identity (**Figure 1E**) [3]. αβT cell receptor (TCR) sequencing was also performed to assess the identity of T cell clones derived from the bronchoalveolar compartment. All clones utilized TRAV1-2/TRAJ33 TCRα chains, consistent with canonical MAIT cell TCRα sequences and all had MAIT match scores of 1.00 based on CDR3α similarity to published MAIT cell receptor sequences (**Figure 1F**).

### Lung mucosal MAIT cells are functionally inhibited compared to peripheral counterparts

Once we had established that CD3+CD4-MR1 tetramer-positive bronchoalveolar T cells with atypical MAIT cell phenotype were MR1-restricted MAIT cells, we sought to understand their functional capacity. Because MAIT cells have been shown to produce Th1 and Th17 cytokines as well as cytolytic products, we assessed the constitutive and stimulated production of IFN-γ, IL-17 and granzyme B in *ex-vivo* samples (characteristic example and gating strategy shown in **Figure 2A**). Although using MAIT-ligand specific stimulations would have been optimal, the *ex-vivo* samples had insufficient cell numbers to support multiple stimulations. Thus, we stimulated with PMA/ionomycin, a non-specific mitogen that has been shown to induce production of IL-17 from MAIT cells [28]. PMA/ionomycin stimulation of peripheral blood MAIT cells from healthy HIV-negative controls resulted in significantly higher IFN-γ production than that seen in matched bronchoalveolar MAIT cells (*P* = 0.0156) (**Figure 2B**). Bronchoalveolar MAIT cells produced significantly less IFN-γ than matched MR1 tetramer-negative CD8+ T cells (**Supplementary Figure 1A**). Both peripheral blood and bronchoalveolar MAIT cells produced low levels of IL-17 upon stimulation (**Figure 2C**), which in the case of peripheral blood MAIT cells was significantly greater than that produced by matched MR1 tetramer-negative CD8+ T cells (**Supplementary Figure 1B**). Constitutive production of granzyme B by peripheral and bronchoalveolar MAIT cells was low in both compartments (Figure 2C) and lower than that produced by matched MR1 tetramer-negative CD8+ T cells (*P* < 0.0001 in both compartments) (**Supplementary Figure 1C**). To determine whether the reduced pro-inflammatory function of bronchoalveolar MAIT cells might be associated with expression of inhibitory co-receptors, we assessed surface expression of PD-1 and TIM-3 on MAIT cells from both compartments. We found both of these markers to be expressed at significantly higher levels in bronchoalveolar MAIT cells compared to their peripheral counterparts (**Figure 2E and F**).

**Figure 2:**
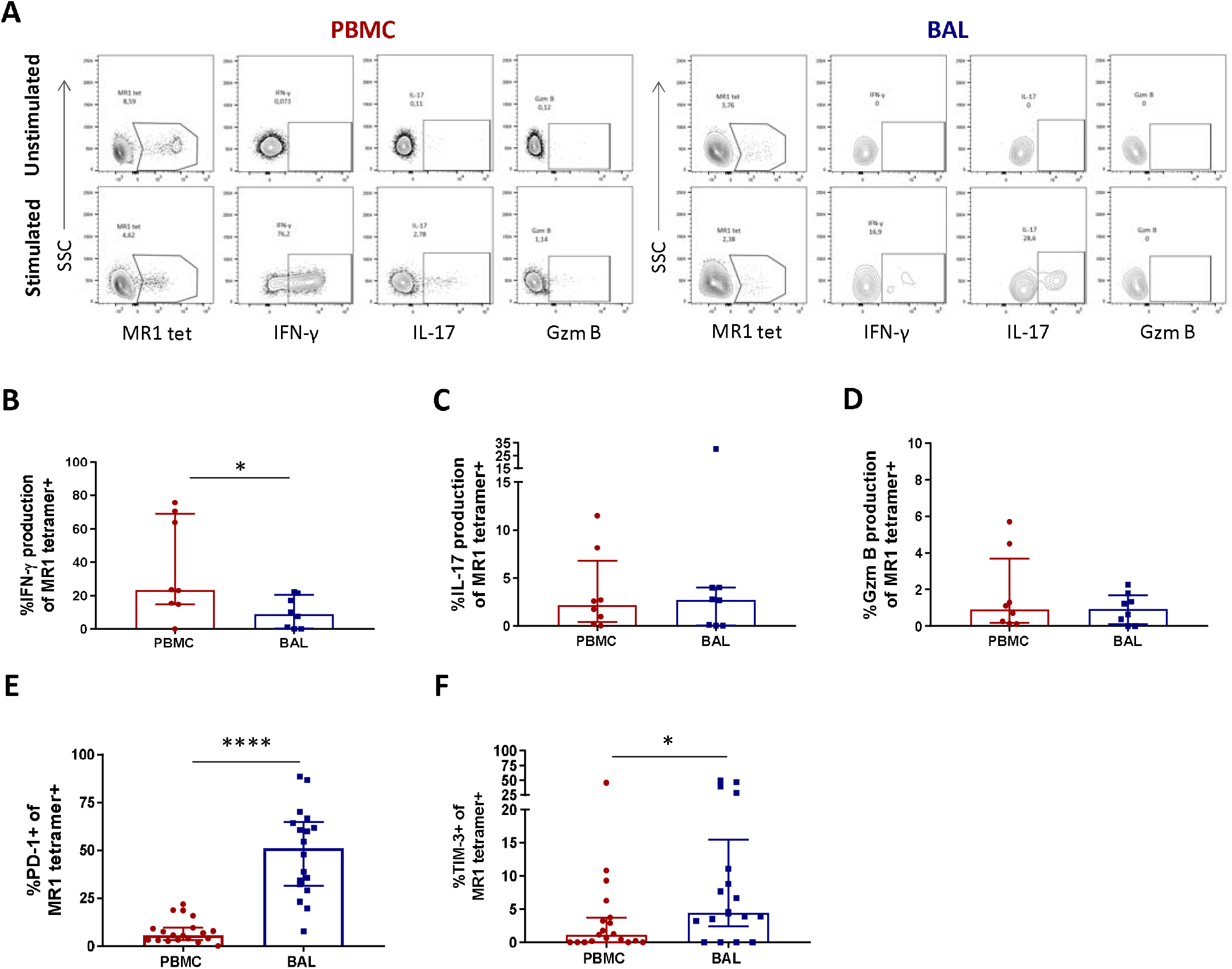
Peripheral blood MAIT cells display a pro-inflammatory profile in healthy participants. (A) Gating strategy used to define cytokine producing CD3+CD4-MR1 5-OP-RU tetramer-positive cells observed in the unstimulated (top row) and stimulated (bottom row) conditions. *Ex vivo* staining of peripheral blood and bronchoalveolar MAIT cells for the production of (B) inducible IFN-γ (*P* = 0.0156), (C) inducible IL-17 and (D) constitutive granzyme B following 6 hour stimulation with PMA/ionomycin (n = 8). Frequency of (E) PD-1 (P < 0.0001) and (F) TIM-3 (*P* = 0.0417) expressing peripheral blood (red) and bronchoalveolar (blue) MAIT cells (n = 18). Bars represent medians and error bars represent interquartile ranges. Statistical difference was determined using the Mann-Whitney U test.

### HIV infection leads to a numeric depletion of bronchoalveolar MAIT cell, but phenotype and function are preserved

We next sought to evaluate the impact of HIV infection on MAIT cell number, phenotype and function in the peripheral blood and bronchoalveolar compartment. We first performed MR1 tetramer staining and confirmed MAIT cell depletion in the peripheral blood with median MR1 tetramer-positive cell frequencies declining from 1.74% (0.74 – 3.53%) of CD3+CD4-T cells to 0.88% (0.16 – 2.49%) (*P* = 0.0349, **Figure 3A**). Assessment of CD161 and CD26 expression on their phenotypic subpopulations revealed that while MAIT cells in the peripheral blood remained dominated by the CD161++CD26++ phenotype, regardless of disease state, HIV caused a small but significant shift in the phenotype of peripheral blood MAIT cells with a decrease in the CD161++CD26++ subpopulation (*P* = 0.0202) from 94.2% (87.9 – 97.9%) to 84.3% (74.83 – 94.43%) and increase in the CD161-CD26-subpopulation (*P* = 0.0104) from 1.35% (0 – 3.73%) to 5.38% (2.48 – 12.95%) (**Figure 3B**). MR1 tetramer staining revealed MAIT cells were numerically depleted in the bronchoalveolar compartment with median MR1 tetramer-positive cell frequencies declining from 1.15% (0.44 −2.22%) to 0.38% (0.21 – 1.58%) (*P* = 0.0471, **Figure 3C**). In the bronchoalveolar compartment, the phenotypic heterogeneity of MAIT cells was maintained during HIV infection with the proportion of MAIT cells with the typical CD161++CD26++ phenotype being reduced from a median of 64.95% (39.0 – 77.88%) to 37.63% (15.68 – 52.7%) (*P* = 0.0176) and the frequencies of the atypical phenotypes non-significantly increased (**Figure 3D**). Analysis of the functional capacity of ex-vivo stimulated MAIT cells showed no significant difference in inducible IFN-γ production in the HIV-positive group in either compartment, although there was a non-significant trend towards a reduction in the peripheral blood of HIV (**Figure 3E**). IL-17 producing MAIT cells were reduced (*P* = 0.0247) in the peripheral blood, but not significantly altered in the bronchoalveolar compartment (**Figure 3F**). HIV infection did not alter constitutive granzyme B production in either the peripheral blood or the bronchoalveolar compartment (**Figure 3G**).

**Figure 3:**
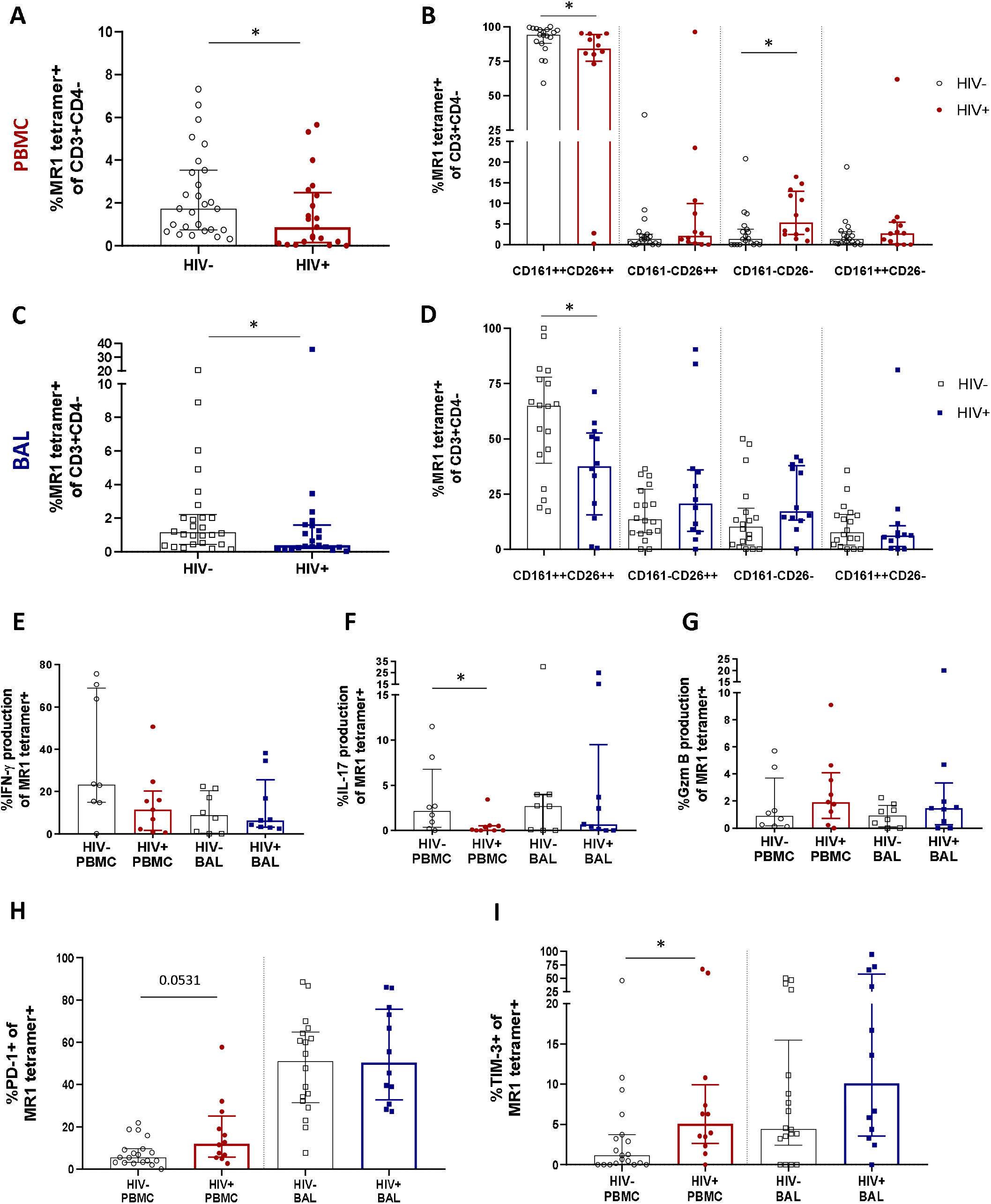
Preservation of bronchoalveolar MAIT cell phenotype and function in HIV infection. (A) MR1 tetramer staining of CD3+CD4-cells showing depletion peripheral blood MAIT cells (*P* = 0.0349) and (B) CD161 and CD26 staining of peripheral blood MAIT cells (*P* = 0.0202 and 0.0104). (C) Depletion of bronchoalveolar MAIT cells (*P* = 0.0471) and (D) CD161 and CD26 staining of bronchoalveolar MAIT cells (*P* = 0.0176) [HIV-negative (n = 18) and HIV-positive (n = 12) participants are shown]. Intracellular cytokine staining for (E) inducible IFN-γ, (F) inducible IL-17 (*P* = 0.0247) and (G) constitutive granzyme B production in HIV-negative (n = 8) and HIV-positive (n = 9) peripheral blood and bronchoalveolar MAIT cells following 6 hour stimulation with PMA/ionomycin. Frequency of (H) PD-1 and (I) TIM-3 (*P* = 0.0171) expressing MR1 tetramer+ cells in the peripheral blood and bronchoalveolar lavage fluid of HIV-negative and HIV-positive individuals. Bars represent medians and error bars represent interquartile ranges. Statistical difference was determined using the Mann-Whitney *U* test.

In HIV-positive participants, PD-1 expression in bronchoalveolar MAIT cells remained unchanged, while a trend towards increased PD-1 expression was observed in peripheral blood (*P* = 0.0531) MAIT cells (**Figure 3H**). Interestingly, a sub-analysis of CD8+ (**Supplementary Figure 2A**) and CD4-CD8-MAIT cells (**Supplementary Figure 2B**) showed that PD-1 expression was significantly increased on the CD8+ MAIT cells (*P* = 0.0019), but not on the CD4-CD8-MAIT cells in the peripheral blood of HIV-positive participants. PD-1 expression remained unchanged in both CD8+ and CD4-CD8-MAIT cells in the bronchoalveolar compartment in HIV. TIM-3 expression showed a trend towards higher expression in both compartments in people with HIV (**Figure 3I**) but was significantly elevated only in peripheral blood MAIT cells (*P* = 0.0171). Overall, these findings suggest that HIV infection reduces the functional capacity and induces expression of inhibitory receptors on peripheral blood MAIT cells but in the bronchoalveolar compartment, where functional capacity is lower and inhibitory markers are higher even in healthy HIV-negative individuals, there is little additional impact of HIV infection on MAIT cell function.

### Transcriptomic heterogeneity of MAIT cells

Having found that bronchoalveolar MAIT cells display greater phenotypic heterogeneity compared to peripheral blood MAIT cells, and that HIV reduces the function and induces inhibition in peripheral MAIT cells while having relatively less impact on the bronchoalveolar compartment, we next sought to better understand the phenotypic heterogeneity of bronchoalveolar MAIT cells using single-cell RNA-sequencing. Unsupervised analysis of 190 MR1 tetramer-positive cells pooled from 9 individuals, including HIV-negative and HIV-positive samples with paired samples from both compartments, revealed that MAIT cells separate into four distinct transcriptional clusters based upon unique gene expression patterns (**Figure 4A**). Clustering was highly driven by compartment (**Figure 4B**) with Cluster_0 (coral) and Cluster_3 (purple) composed predominantly of bronchoalveolar MAIT cells (75.44% and 69.70% respectively, **Supplementary Figure 3A**) and thus we renamed them BAL_1 and BAL_2; meanwhile, the remaining 2 clusters (Cluster_1 (lime green) and Cluster_2 (blue)) were mostly made up of peripheral blood MAIT cells (86.54% and 64.58% respectively) and thus we renamed them PBMC_1 and PBMC_2. HIV status did not appear to be a driver of these unsupervised clusters with cells from HIV-negative and HIV-positive participants represented in each cluster (**Supplementary Figures 3A and B**). Clusters were also represented by cells from various individuals, with inter-individual differences noted in the distribution of the transcriptional subsets (**Supplementary Figure 4**).

**Figure 4:**
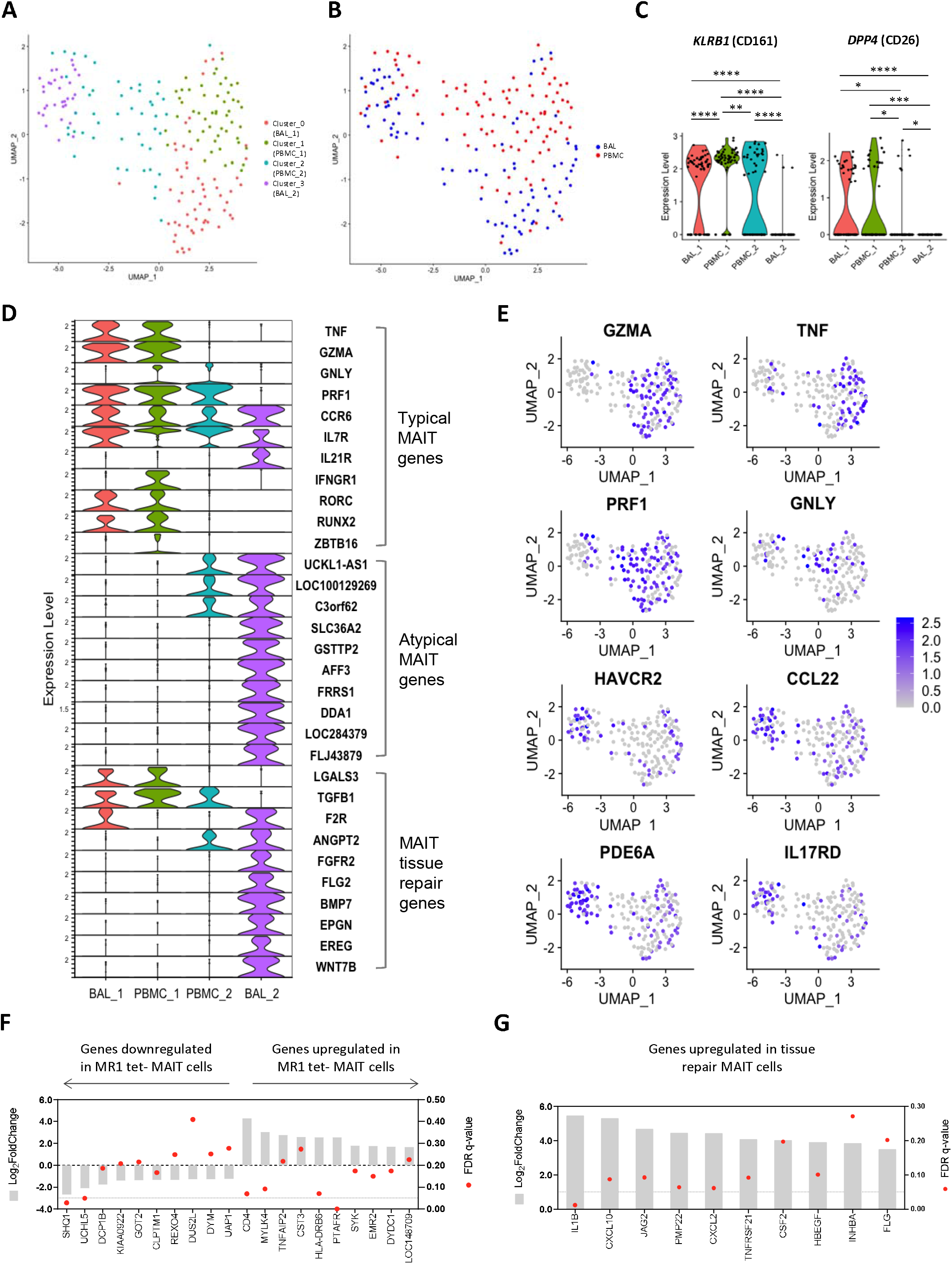
Transcriptomic heterogeneity of MAIT cells. (A) UMAP plot showing clustering of MR1 tetramer+ MAIT cells into four distinct transcriptomic subsets by unsupervised analysis. (B) UMAP plot showing assignment of compartment of origin to MAIT cell transcriptomic subsets. (C) Violin plot showing the expression of MAIT cell markers *KLRB1* (CD161) and *DPP4* (CD26) by each transcriptomic subset. (D) Violin plot showing the expression of typical MAIT cell genes (top), atypical MR1 tetramer-negative MAIT cell genes (middle), and tissue repair genes (bottom) by MR1 tetramer-positive MAIT cells from each transcriptomic subset. (E) UMAP plots showing genes of interest expressed by BAL_2 MAIT cells, including MAIT effector genes, atypical MR1 tetramer-negative MAIT genes and tissue repair genes. Bulk RNA-sequence analysis showing (F) the enrichment of atypical MAIT gene signature and (G) the upregulation of MAIT tissue repair genes in the bronchoalveolar MAIT cells of healthy individuals as compared to peripheral blood MAIT cells.

The unsupervised transcriptomic clusters demonstrated interesting patterns of differential expression of the two typical MAIT cell genes, *KLRB1* (CD161) and *DPP4* (CD26) (**Figure 4C**). The BAL_1 and PBMC_1 subsets were characterized by transcriptional co-expression *KLRB1* and *DPP4*, while the PBMC_2 subset expressed *KLRB1* alone and the BAL_2 subset expressed neither. We assessed the expression levels of other genes known to be characteristic of the MAIT cell transcriptome, termed “typical” MAIT genes, including effector molecules, cytokine receptors and transcription factor transcripts in the four MAIT cell subsets[9, 29–32]. Again, we found generally high levels of expression of typical MAIT genes in the BAL_1 and PBMC_1 subsets with a few of these also expressed in the PBMC_2 subset (**Figure 4D and 4E**). Of the representative typical MAIT cell transcripts, only a few (*CCR6*, *IL7R* and *IL21R*) were expressed by the BAL_2 subset. Other transcripts in the typical MAIT gene category were absent or expressed at very low levels in the BAL_2 MAIT subset. We first considered the possibility that our MR1 tetramer-positive cells had been contaminated with alveolar macrophages, a highly prevalent population in BAL fluid, by determining the expression of known macrophage genes in this cluster such as *MARCO, VSIG4 and CD14*. We found there to be very little expression of these transcripts (**Supplementary Figure 5**) and concluded that the BAL_2 subset did not consist of macrophage contaminants. Having found phenotypically atypical MR1 tetramer-positive cells in the bronchoalveolar compartment, which we confirmed to be MR1-restricted MAIT cells by functional assay, we hypothesized that BAL_2 may be composed of atypical MAIT cells. Next, we assessed the expression of genes found to be elevated in a population of recently described atypical MR1 tetramer-negative MAIT cells [9] and found that these genes, which we termed “atypical MAIT genes” (listed in **Supplementary Table 1**) were more frequently expressed in BAL_2 than any of the other MAIT cell transcriptional subsets (**Figure 4D** and *PDE6A* and *IL17RD* in **Figure 4E**), suggesting a resemblance to these novel MAIT cells with atypical gene expression. Interestingly, the PBMC_2 subset appeared to have an intermediate transcriptional character, expressing some but not all of the typical MAIT cell genes and a few of the atypical MAIT cell genes. In addition to having pro-inflammatory and cytotoxic functions, MAIT cells have been recently shown to also be involved in tissue repair [33–35]. By combining genes lists obtained from literature [33–35], we compiled a list of 131 genes expressed in MAIT cells associated with tissue repair functions (**Supplementary Table 2**). Interestingly, we found that while a few of these genes that were detectable in our single-cell dataset were present in BAL_1, PBMC_1 and PBMC_2 (*LGALS3* and *TGFB1*), the majority of these were exclusively upregulated in the BAL_2 cluster (**Figure 4D**). Because our protein-level data had shown that bronchoalveolar MAIT cells express more PD-1 and TIM-3, we next assessed the transcriptomic expression of these inhibitory markers and found that cells in the BAL_2 subset displayed greater expression of *HAVCR2* (TIM-3) (**Figure 4E**), expression of which may contribute to the low expression of effector gene transcripts that we observed in this subset. No differences were noted in *PDCD1* (PD-1) expression, which was relatively low across all transcriptomic subsets, potentially due to limitations of single-cell sequencing. Taken together these results suggest that three of our unsupervised clusters (BAL_1, PBMC_1 and PBMC_2) share expression of many genes that are known to be typical of MAIT cells, but that the fourth cluster (BAL_2) was characterized by distinct gene expression that shared similarities with two groups of recently published alternative MAIT cell phenotypes. Additionally, our data suggest that the peripheral blood is composed of MAIT cell subsets with subtle transcriptomic differences (PBMC_1 and PBMC_2) while the bronchoalveolar compartment consists of two transcriptionally distinct subsets (BAL_1 and BAL_2).

Because single-cell transcriptional data has limitations, and the detection of certain important transcripts in our single-cell dataset was very low, we next used bulk RNA-sequencing obtained from 100 cell mini-populations of MR1 tetramer-positive cells to ascertain whether alternative MAIT cell gene signatures were found to be more highly expressed in bronchoalveolar MR1 tetramer-positive cells compared to their peripheral counterparts in healthy HIV-negative people. To assess for the expression of the atypical MAIT cell genes, we used a list of 217 differentially expressed genes (51 downregulated and 166 upregulated genes) from Pomaznoy *et al.* [9] (**Supplementary Table 1**), and found that 82% of the genes that were downregulated by atypical MAIT cells in the original dataset were downregulated in the bronchoalveolar MR1 tetramer-positive cells in our dataset and that 62% of the genes upregulated by atypical MAIT cells in their dataset were upregulated in our dataset (top 10 down- and upregulated genes shown in **Figure 4F**). This finding suggests an enrichment of atypical MAIT cells in the bronchoalveolar compartment. Using the list of 131 genes that are upregulated in tissue repair MAIT cells, we similarly found that 67% of these tissue repair genes were upregulated in bronchoalveolar MAIT cells (**Figure 4G** and **Supplementary Table 2**). Together this bulk RNA transcriptional analysis supported our conclusion that bronchoalveolar MR1 tetramer-positive cells may comprise two distinct subsets: one of which has transcriptional features typical of peripheral MAIT cells and the other which shares transcriptional features with MR1 tetramer-negative MAIT cells and may possess tissue repair functions.

## Discussion

Previous work detailing HIV-induced alterations of MAIT cells largely focused on peripheral blood responses despite there being evidence that MAIT cells may be important in the response to infection at mucosal surfaces. In this study, we aimed to address this gap in knowledge by examining MAIT cell phenotype, function and transcriptome in the lung mucosa of healthy humans and determining how these features are altered in people with HIV infection. We found that bronchoalveolar MAIT cells in healthy individuals feature previously undescribed phenotypic heterogeneity and were less pro-inflammatory, displaying greater expression of inhibitory co-receptors, than their peripheral blood counterparts. Our analysis of MAIT cells in people living with HIV showed that while HIV decreases the frequency of MAIT cells at both anatomical sites, its abrogation of MAIT cell function is more marked in the peripheral blood than at the lung mucosal surface. Using single-cell RNA-sequencing we were able to uncover the presence of two distinct MAIT cell subsets in the bronchoalveolar compartment, which would have been overlooked had we exclusively used bulk RNA-sequencing. The first bronchoalveolar subset shares the well-described transcriptional features of typical peripheral blood MAIT cells while the second has few of the typical MAIT cell transcriptional features and instead expresses transcriptional programs associated with atypical MR1 tetramer-negative MAIT cells and alternative MAIT tissue repair functions. The phenotypic and transcriptional heterogeneity we observed in bronchoalveolar MAIT cells was preserved even in people who are HIV-positive.

We found that bronchoalveolar MAIT cells of healthy humans feature previously unreported phenotypic heterogeneity with regard to CD161 and CD26 expression. The expression of both CD161 and CD26 is associated with cytotoxicity in NK cells [36], with regulatory T (Treg) cells being characterised by low/no expression of CD26 [37]. In addition to being more phenotypically heterogenous, bronchoalveolar MAIT cells had a less pro-inflammatory phenotype than their peripheral blood counterparts, producing less inducible IFN-γ upon PMA/ionomycin stimulation. The pro-inflammatory capacity of peripheral blood MAIT cells via the production of IFN-γ and TNF-α has been well documented [3, 5]. MAIT cells from the lung’s mucosal surface during active TB have been shown to have higher pro-inflammatory function (specifically TNF-α production) than peripheral counterparts [7]. Lower pro-inflammatory responses have been shown in both nasopharyngeal and oral mucosal MAIT cells, with the latter being paired with increases in the expression of HLA-DR and PD-1 [3, 5]. Thus, the higher expression of inhibitory markers we observed in bronchoalveolar MAIT cells may not only function to minimize exaggerated inflammatory responses to commensal microbes at the lung mucosa, but paired with the downregulation of CD161 and CD26, may also be indicative of alternative functions of these MAIT cells, not relating to cytotoxicity.

Characterization of MAIT cell phenotype and function in HIV-positive individuals showed a decline in peripheral blood MAIT cells in HIV infection, as has been previously reported [16, 17], as well as a decline in MAIT cell frequency in the bronchoalveolar compartment. This finding is consistent with previous work that has reported a decline in lung mucosal MAIT cells in humans using the phenotypic definition of MAIT cells (CD8+CD161++TRAV1-2+), where MAIT cell frequency was reduced in a manner that was inversely correlated to viral load [18], as well as in the rhesus macaque model of SIV infection by MR1 tetramer definition [38]. We found that MAIT cell surface marker heterogeneity was preserved in bronchoalveolar MAIT cells in HIV infection, however, we did note a reduction in the frequency of CD161++CD26++ MAIT cells in both the bronchoalveolar and peripheral blood compartments. In the peripheral blood this was associated with an increase in the frequency of CD161-CD26-MAIT cells. Leeansyah *et al.* [17] and Eberhard *et al.* [27] both reported an increase in CD161-negative MAIT cells following the reduction in CD161-positive MAIT cells attributed to MAIT cell activation due to persistent antigen exposure in HIV infected individuals, which may also be the case in our setting. In HIV-positive individuals, MAIT cell pro-inflammatory capacity was reduced in peripheral blood MAIT cells, as demonstrated by the decreased ability to produce IL-17, and higher TIM-3 surface expression. Our finding of increased expression of inhibitory receptors in peripheral blood MAIT cells of HIV-positive people may explain the decline in peripheral blood MAIT cell function. Bronchoalveolar MAIT cells on the other hand, displayed no significant alterations in function, which contrasts to what has been observed in other T cell subsets including bronchoalveolar CD4+ T cells which are functionally impaired during HIV infection [39].

Transcriptomic characterization of MAIT cells revealed the presence of a distinct subset occupying the bronchoalveolar compartment, BAL_2, which was found to resemble the recently described MR1 tetramer-negative MAIT cells which were enriched in latently TB infected individuals and had greater TCR β-chain diversity suggesting an increased capacity for antigen discrimination [9]. Consistent with the hypothesis of less cytotoxic function in cells where *KLRB1* (CD161) and *DDP4* (CD26) expression is low or absent, we found cells in the BAL_2 subset devoid of the expression of typical MAIT effector genes such as *GZMA, TNF, GNLY* and *PRF1*, but instead found that they expressed atypical genes such as *PDE6A* and *IL17RD* and tissue repair genes like *CCL22*. Not only is *CCL22* involved in tissue repair [33–35], but together with IL17RD may also play an immunoregulatory role. The expression of CCL22 promotes interactions between dendritic cells (DCs) and Tregs which then leads to the dampening of immune responses [40], while IL17RD reduces pro-inflammatory signalling via TLRs as well as the expression of pro-inflammatory genes like *IL-6* [41, 42]. MAIT cells belonging to the BAL_2 subset also displayed high expression of typical MAIT gene *IL21R* in contrast to MAIT cells in other transcriptional subsets as well as *HAVCR2* (TIM-3). The expression of IL-21R on T cells is associated with maintained function during chronic infection [43], with IL-21R^−/−^ mice having increased susceptibility to mycobacterial infection [44]. Exposure of lung mucosal MAIT cells to greater microbial diversity in the respiratory tract, may lead to the expansion of a subset of MAIT cells with atypical features and greater ligand discrimination, similar to the MR1 tetramer-negative MAIT cells observed in the peripheral blood of people with latent TB [9]. Alternatively, MAIT cell heterogeneity may occur as a result to exposure to non-microbial environmental cues at the lung mucosa. Atypical and MAIT tissue repair transcripts were also enriched in bulk-sorted bronchoalveolar MAIT cell populations, likely because bulk-sorting combined the two BAL-resident transcriptional subpopulations of MAIT cells. In contrast to typical pro-inflammatory MAIT cell functions, the BAL_2 subset specifically may have typical effector functions inhibited by high expression of *HAVCR2* (TIM-3) and play an alternative function contributing to the prevention of excessive tissue damage in the bronchoalveolar compartment.

Our study had a number of limitations. Our functional assays were limited by the number of available cells and thus utilized non-specific stimuli and were performed on only a subset of the cohort. Certain important genes were not well-represented in our single-cell dataset, limiting our ability to fully determine the transcriptional characteristics of the novel, BAL_2 subset. Use of MR1 5-OP-RU tetramer may have neglected to identify MAIT cells that respond to alternate small molecule antigens [6]. In addition, all HIV-positive participants were antiretroviral therapy naive, leaving important questions about MAIT cells in people on effective ART unanswered.

In conclusion, we report previously unrecognized phenotypic and transcriptional heterogeneity of MR1 tetramer positive MAIT cells in the bronchoalveolar compartment. Single-cell transcriptional analysis suggests the existence of a distinct subset of MAIT cells at the lung mucosal surface that expresses low levels of most typical MAIT cell effector genes and is characterized by expression of alternative tissue repair and inhibitory genes. Further research is required to determine if these cells possess an immunoregulatory role and if they contribute to the maintenance of homeostasis in the lung environment. We found that HIV reduced the number of MAIT cells in both peripheral blood and bronchoalveolar compartments and confirmed previous findings that HIV infection induces expression of inhibitory receptors and reduces the functional capacity of circulating MAIT cells. In contrast, at the lung’s mucosal surface where MAIT cell functional capacity is lower and inhibitory markers are higher even in healthy HIV-negative individuals, we found little additional impact of HIV infection on MAIT cell function. Finally, our data suggest the relative preservation of MAIT cell phenotypic and transcriptional heterogeneity at the lung’s mucosal surface even in the face of untreated HIV infection.

## Acknowledgments

We would like to thank the study participants and the clinical staff of Inkosi Albert Luthuli Hospital, without whom this work would not be possible. We would also like to thank the AHRI Immunology core, the OHSU Flow Cytometry core, the AHRI Biorepository, Clinical and Microbiology cores. The MR1 tetramer technology was developed jointly by Dr James McCluskey, Dr Jamie Rossjohn, and Dr David Fairlie, and the material was produced by the NIH Tetramer Core Facility as permitted to be distributed by the University of Melbourne.

## Data availability statement

All source data used for transcriptomic analyses are available at Figshare: 10.6084/m9.figshare.13259657

## Author contributions

SK, MMt, EWM, SMP and TB performed the experiments; SK, MMt, EWM, SMP, SWK and EBW analysed the results; KN, DFK, PM, MMi, MS, ZM, DR, and FK enrolled participants, collected samples and performed clinical procedures. AKS, DML, TN and EBW provided supervision; SK and EBW wrote the manuscript. All co-authors reviewed and approved the final manuscript.

## Supplementary Table and Supplementary Figure Legends

**Supplementary Table 1**: Genes differentially expressed by atypical MR1 tetramer-negative MAIT cells during latent TB infection and similarly differentially expressed by bronchoalveolar MAIT cells as compared to peripheral blood MAIT cells.

**Supplementary Table 2**: MAIT tissue repair genes also upregulated by bronchoalveolar MAIT cells in comparison to peripheral blood MAIT cells.

**Supplementary Figure 1**: Pro-inflammatory cytokine and cytolytic molecule production of MR1 tetramer-positive MAIT cells (n = 8) in the peripheral blood (red) and bronchoalveolar lavage (BAL) fluid (blue) in contrast to matched conventional CD8+ T cells (n = 10) from healthy participants showing (A) inducible IFN-γ, (B) inducible IL-17 and (C) constitutive granzyme B production. Bars represent medians and error bars represent interquartile ranges. Statistical difference was determined using the Mann-Whitney *U* test.

**Supplementary Figure 2**: Frequency of PD-1 expressing (A) CD8+ MR1 tetramer-positive MAIT cells and (B) CD4-CD8-MR1 tetramer-positive MAIT cells from the peripheral blood of HIV-negative and HIV-positive participants. Bars represent medians and error bars represent interquartile ranges. Statistical difference was determined using the Mann-Whitney *U* test.

**Supplementary Figure 3**: (A) Summary of characteristics of MR1 tetramer-positive MAIT cell transcriptomic clusters. (B) UMAP plot showing assignment of HIV status to MAIT cell transcriptomic subsets.

**Supplementary Figure 4**: Percentage distribution of MAIT cell transcriptomic subsets by each participant identifier (PID).

**Supplementary Figure 5**: Heatmap showing the expression of canonical macrophage genes across the four transcriptional MAIT cell subsets.

