## Supplementary Tables for "MR1-restricted MAIT cells from the human lung mucosal surface have distinct phenotypic, functional, and transcriptomic features that are preserved in HIV infection"

**Supplementary Table 1:** Genes differentially expressed by atypical MR1 tetramer‑negative MAIT cells during latent TB infection and similarly differentially expressed by bronchoalveolar MAIT cells as compared to peripheral blood MAIT cells.

| Downregulated genes | | | Upregulated genes | | |
| --- | --- | --- | --- | --- | --- |
| Gene | Log_2_Fold Change | Adjusted *P* | Gene | Log_2_Fold Change | Adjusted *P* |
| *SHQ1* | -2.6701 | 0.0284 | *CD4* | 4.2899 | 0.0694 |
| *UCHL5* | -2.1101 | 0.0491 | *MYLK4* | 3.0350 | 0.0921 |
| *DCP1B* | -1.7564 | 0.1866 | *TNFAIP2* | 2.7483 | 0.2181 |
| *KIAA0922* | -1.3915 | 0.2070 | *CST3* | 2.5643 | 0.2736 |
| *GOT2* | -1.3571 | 0.2150 | *HLA-DRB6* | 2.5482 | 0.0698 |
| *CLPTM1* | -1.3420 | 0.1664 | *PTAFR* | 2.5395 | 0.0009 |
| *REXO4* | -1.3369 | 0.2481 | *SYK* | 1.7857 | 0.1743 |
| *DUS2L* | -1.2888 | 0.4092 | *EMR2* | 1.7575 | 0.1492 |
| *DYM* | -1.2671 | 0.2520 | *DYDC1* | 1.6938 | 0.1745 |
| *UAP1* | -1.2588 | 0.2778 | *LOC148709* | 1.6510 | 0.2260 |
| *ADK* | -1.2569 | 0.2680 | *UCKL1-AS1* | 1.2333 | 0.1327 |
| *SF3B4* | -1.2379 | 0.2404 | *FFAR2* | 1.2208 | 0.1693 |
| *G6PD* | -1.1753 | 0.2812 | *TMEM170B* | 1.1443 | 0.1714 |
| *SUCLG2* | -1.1519 | 0.3105 | *LOC284379* | 1.1325 | 0.1008 |
| *TBC1D14* | -1.0872 | 0.3873 | *POU5F1* | 1.1270 | 0.1433 |
| *EHMT1* | -1.0569 | 0.2885 | *PARD6G* | 1.1220 | 0.0832 |
| *C17orf62* | -1.0178 | 0.3102 | *C4orf26* | 1.1031 | 0.1397 |
| *DDB1* | -1.0110 | 0.3546 | *INMT* | 1.0740 | 0.1059 |
| *PSMC4* | -0.9982 | 0.3227 | *NLRP12* | 1.0387 | 0.1513 |
| *MDH2* | -0.9210 | 0.3924 | *ITGA2* | 1.0100 | 0.2177 |
| *FNTA* | -0.8901 | 0.4044 | *SLC16A12* | 0.9879 | 0.3541 |
| *NFYB* | -0.8863 | 0.3046 | *LOC100506385* | 0.9696 | 0.2255 |
| *P4HTM* | -0.8772 | 0.4338 | *CNNM1* | 0.9499 | 0.2671 |
| *SAE1* | -0.8015 | 0.4571 | *BHMT2* | 0.9432 | 0.1588 |
| *RHBDD2* | -0.7844 | 0.4511 | *MBOAT2* | 0.9405 | 0.2610 |
| *PSMC2* | -0.7532 | 0.4045 | *NDST3* | 0.8965 | 0.2561 |
| *DENND2D* | -0.6513 | 0.5297 | *C9orf66* | 0.8714 | 0.2814 |
| *KCNA3* | -0.6052 | 0.5338 | *IL17RD* | 0.8516 | 0.2422 |
| *CASP8* | -0.5869 | 0.2509 | *GSTTP2* | 0.8512 | 0.3037 |
| *AP3S2* | -0.5843 | 0.4798 | *FBLIM1* | 0.8419 | 0.1897 |
| *KDELR2* | -0.5626 | 0.5786 | *LOC100292680* | 0.8375 | 0.5657 |
| *YTHDF2* | -0.5545 | 0.5784 | *FOXP4* | 0.8366 | 0.2492 |
| *ARCN1* | -0.4939 | 0.4761 | *ADCY1* | 0.8257 | 0.3401 |
| *ATF2* | -0.4807 | 0.6711 | *PART1* | 0.8200 | 0.2501 |
| *HADHB* | -0.4558 | 0.6642 | *EMX2OS* | 0.8016 | 0.2076 |
| *SSR1* | -0.3165 | 0.4241 | *TMEM17* | 0.7826 | 0.3050 |
| *CCR1* | -0.3157 | 0.8277 | *IAPP* | 0.7728 | 0.2636 |
| *ACTR3* | -0.1953 | 0.8609 | *C21orf62* | 0.7703 | 0.1990 |
| *ACAD11* | -0.1938 | 0.8798 | *LOC100129269* | 0.7659 | 0.2206 |
| *CNDP2* | -0.1604 | 0.8786 | *S1PR3* | 0.7407 | 0.4457 |
| *PTPN22* | -0.1474 | 0.8947 | *C14orf105* | 0.7352 | 0.3690 |
| - | - | - | *RAB3B* | 0.7261 | 0.2027 |
| - | - | - | *PTK6* | 0.7249 | 0.2904 |
| - | - | - | *MSRB3* | 0.7112 | 0.2961 |
| - | - | - | *C1orf140* | 0.7106 | 0.3481 |
| - | - | - | *LOC100128338* | 0.7095 | 0.2399 |
| - | - | - | *FRRS1* | 0.7081 | 0.3074 |
| - | - | - | *ST6GAL2* | 0.7039 | 0.3916 |
| - | - | - | *FCAR* | 0.7026 | 0.2764 |
| - | - | - | *PSME4* | 0.6888 | 0.4503 |
| - | - | - | *AGMO* | 0.6763 | 0.4114 |
| - | - | - | *OLFML2A* | 0.6762 | 0.2584 |
| - | - | - | *CEACAM8* | 0.6610 | 0.4789 |
| - | - | - | *KREMEN1* | 0.6568 | 0.2485 |
| - | - | - | *LOC100287792* | 0.6423 | 0.2394 |
| - | - | - | *SYNPO2* | 0.6343 | 0.3566 |
| - | - | - | *LAMC2* | 0.6269 | 0.5295 |
| - | - | - | *TLCD2* | 0.5985 | 0.2482 |
| - | - | - | *CABP4* | 0.5924 | 0.2043 |
| - | - | - | *IRGQ* | 0.5852 | 0.1588 |
| - | - | - | *TTL* | 0.5747 | 0.6595 |
| - | - | - | *VSTM4* | 0.5738 | 0.3236 |
| - | - | - | *TRIM58* | 0.5723 | 0.2671 |
| - | - | - | *PDE6A* | 0.5612 | 0.3774 |
| - | - | - | *LOC100128682* | 0.5426 | 0.2764 |
| - | - | - | *LOC729603* | 0.5388 | 0.3410 |
| - | - | - | *CACNG8* | 0.5231 | 0.2724 |
| - | - | - | *EMP2* | 0.5182 | 0.3263 |
| - | - | - | *CEACAM22P* | 0.4735 | 0.3803 |
| - | - | - | *MYLK3* | 0.4354 | 0.3615 |
| - | - | - | *FLJ43879* | 0.4261 | 0.4326 |
| - | - | - | *LOC284950* | 0.4236 | 0.4711 |
| - | - | - | *CHST6* | 0.3997 | 0.4104 |
| - | - | - | *SLC36A2* | 0.3795 | 0.5738 |
| - | - | - | *POM121L10P* | 0.3724 | 0.4466 |
| - | - | - | *LOC100128288* | 0.3585 | 0.5745 |
| - | - | - | *KLB* | 0.3401 | 0.5633 |
| - | - | - | *NWD1* | 0.3397 | 0.5703 |
| - | - | - | *SLC15A2* | 0.2983 | 0.6820 |
| - | - | - | *FKBP9* | 0.2884 | 0.6537 |
| - | - | - | *ARGFX* | 0.2784 | 0.4804 |
| - | - | - | *CPA4* | 0.2259 | 0.7484 |
| - | - | - | *NAPSB* | 0.2116 | 0.9262 |
| - | - | - | *TSIX* | 0.2041 | 0.7849 |
| - | - | - | *ARSD* | 0.1710 | 0.8481 |
| - | - | - | *C3orf62* | 0.1082 | 0.8512 |
| - | - | - | *LOC400548* | 0.0885 | 0.9055 |
| - | - | - | *FHDC1* | 0.0711 | 0.9386 |
| - | - | - | *LOC286437* | 0.0688 | 0.9345 |
| - | - | - | *AFF3* | 0.0464 | 0.9434 |
| - | - | - | *S100PBP* | 0.0087 | 0.9931 |

**Supplementary Table 2:** MAIT tissue repair genes also upregulated by bronchoalveolar MAIT cells in comparison to peripheral blood MAIT cells.

| Gene | Log_2_Fold Change | Adjusted *P* |
| --- | --- | --- |
| *IL1B* | 5.4659 | 0.0119 |
| *CXCL10* | 5.3120 | 0.0871 |
| *JAG2* | 4.6732 | 0.0926 |
| *PMP22* | 4.4515 | 0.0644 |
| *CXCL2* | 4.4232 | 0.0616 |
| *TNFRSF21* | 4.0764 | 0.0918 |
| *CSF2* | 4.0119 | 0.1972 |
| *HBEGF* | 3.9078 | 0.1008 |
| *INHBA* | 3.8365 | 0.2714 |
| *FLG* | 3.5022 | 0.2018 |
| *APOE* | 3.3867 | 0.2262 |
| *WNT10A* | 3.1996 | 0.2962 |
| *ADM* | 2.7396 | 0.5015 |
| *ZBTB7C* | 2.3640 | 0.4231 |
| *ENG* | 2.0479 | 0.1762 |
| *LGALS3* | 2.0072 | 0.0746 |
| *ADAMTS2* | 1.7944 | 0.2769 |
| *SYK* | 1.7857 | 0.1743 |
| *CXCL12* | 1.5037 | 0.6055 |
| *IGF1* | 1.2513 | 0.1127 |
| *CSF1R* | 1.2232 | 0.6028 |
| *PDGFA* | 1.0010 | 0.6487 |
| *THBS1* | 0.9028 | 0.5151 |
| *FGFR2* | 0.8919 | 0.2309 |
| *CSF1* | 0.7786 | 0.5034 |
| *CCL3* | 0.7190 | 0.6362 |
| *BMP7* | 0.6525 | 0.3417 |
| *DISP1* | 0.5697 | 0.7446 |
| *EREG* | 0.5116 | 0.5041 |
| *FLG2* | 0.4911 | 0.6169 |
| *LEP* | 0.4810 | 0.7144 |
| *EPGN* | 0.4482 | 0.5569 |
| *WNT7B* | 0.3903 | 0.6004 |
| *APP* | 0.3797 | 0.8234 |
| *ANGPT2* | 0.2230 | 0.7109 |
| *CRISPLD2* | 0.1924 | 0.8829 |
| *HIF1A* | 0.1474 | 0.9027 |
| *IFT172* | 0.0849 | 0.9665 |
| *VEGFB* | 0.0816 | 0.9475 |
