## Supplementary Figures for "MR1-restricted MAIT cells from the human lung mucosal surface have distinct phenotypic, functional, and transcriptomic features that are preserved in HIV infection"

### Slide 1
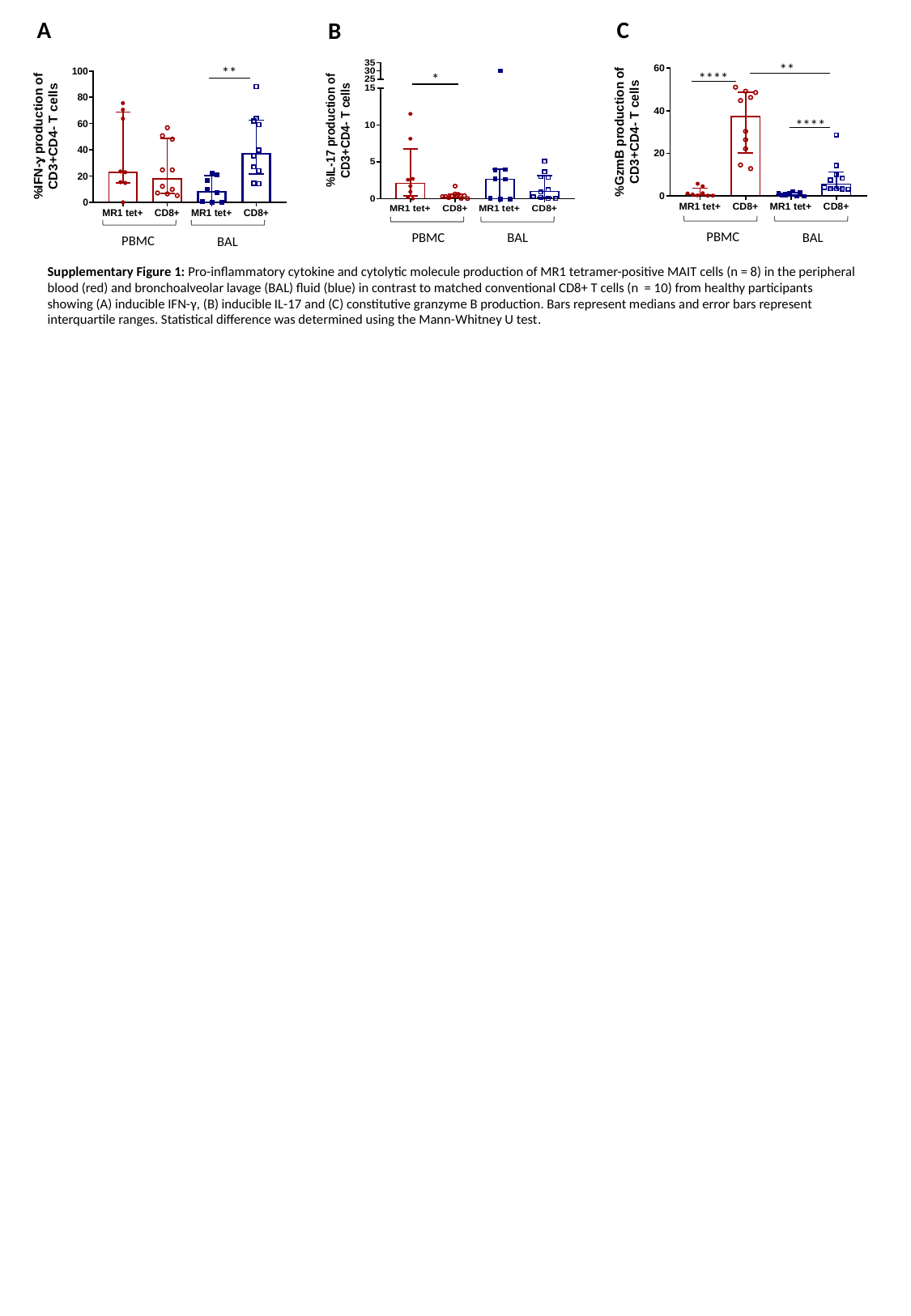

A
C
B
*
BAL
PBMC
**
****
****
PBMC
BAL
**
PBMC
BAL
Supplementary Figure 1: Pro-inflammatory cytokine and cytolytic molecule production of MR1 tetramer-positive MAIT cells (n = 8) in the peripheral blood (red) and bronchoalveolar lavage (BAL) fluid (blue) in contrast to matched conventional CD8+ T cells (n = 10) from healthy participants showing (A) inducible IFN-γ, (B) inducible IL-17 and (C) constitutive granzyme B production. Bars represent medians and error bars represent interquartile ranges. Statistical difference was determined using the Mann-Whitney U test.

### Slide 2
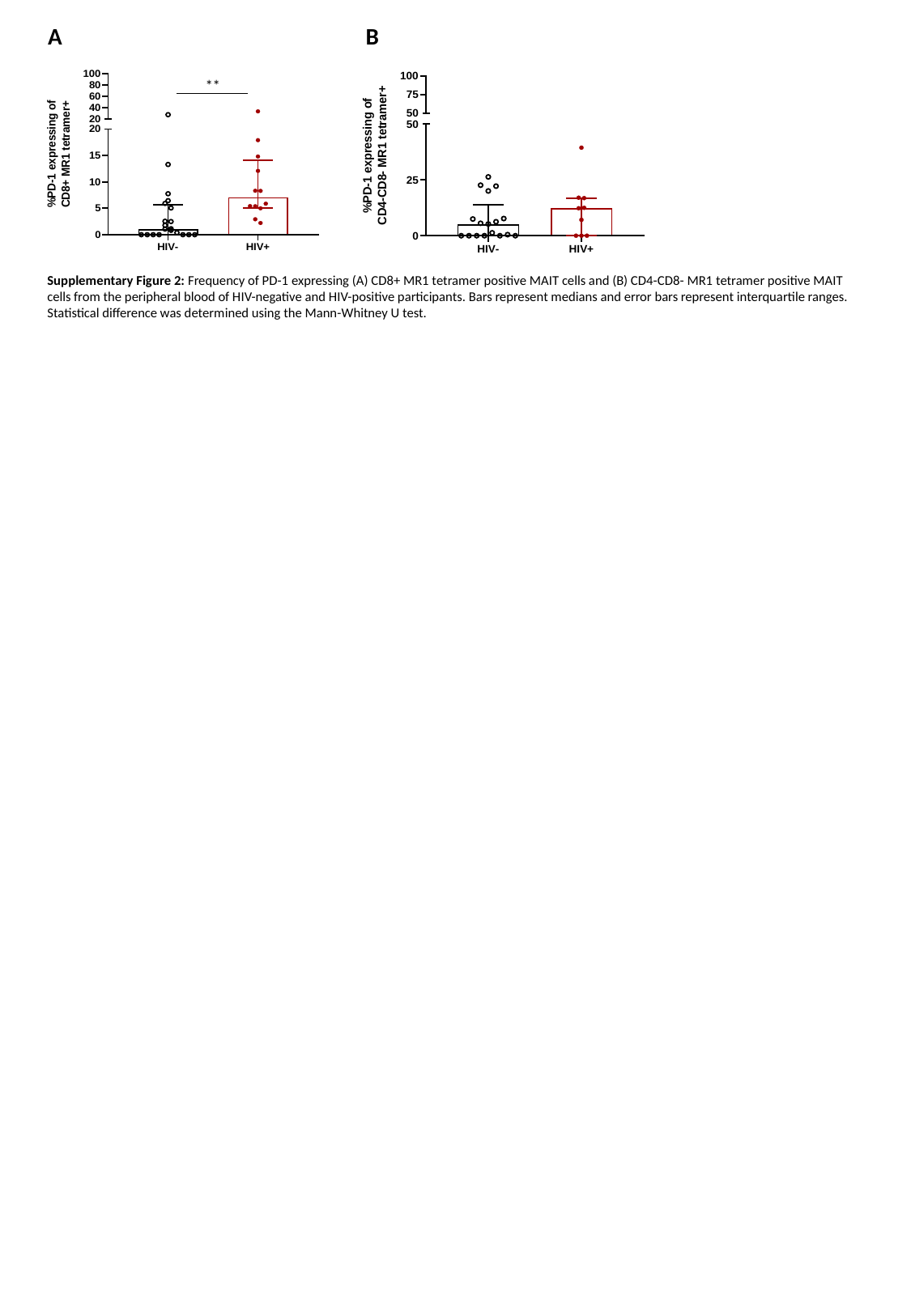

A
B
**
Supplementary Figure 2: Frequency of PD-1 expressing (A) CD8+ MR1 tetramer positive MAIT cells and (B) CD4-CD8- MR1 tetramer positive MAIT cells from the peripheral blood of HIV-negative and HIV-positive participants. Bars represent medians and error bars represent interquartile ranges. Statistical difference was determined using the Mann-Whitney U test.

### Slide 3
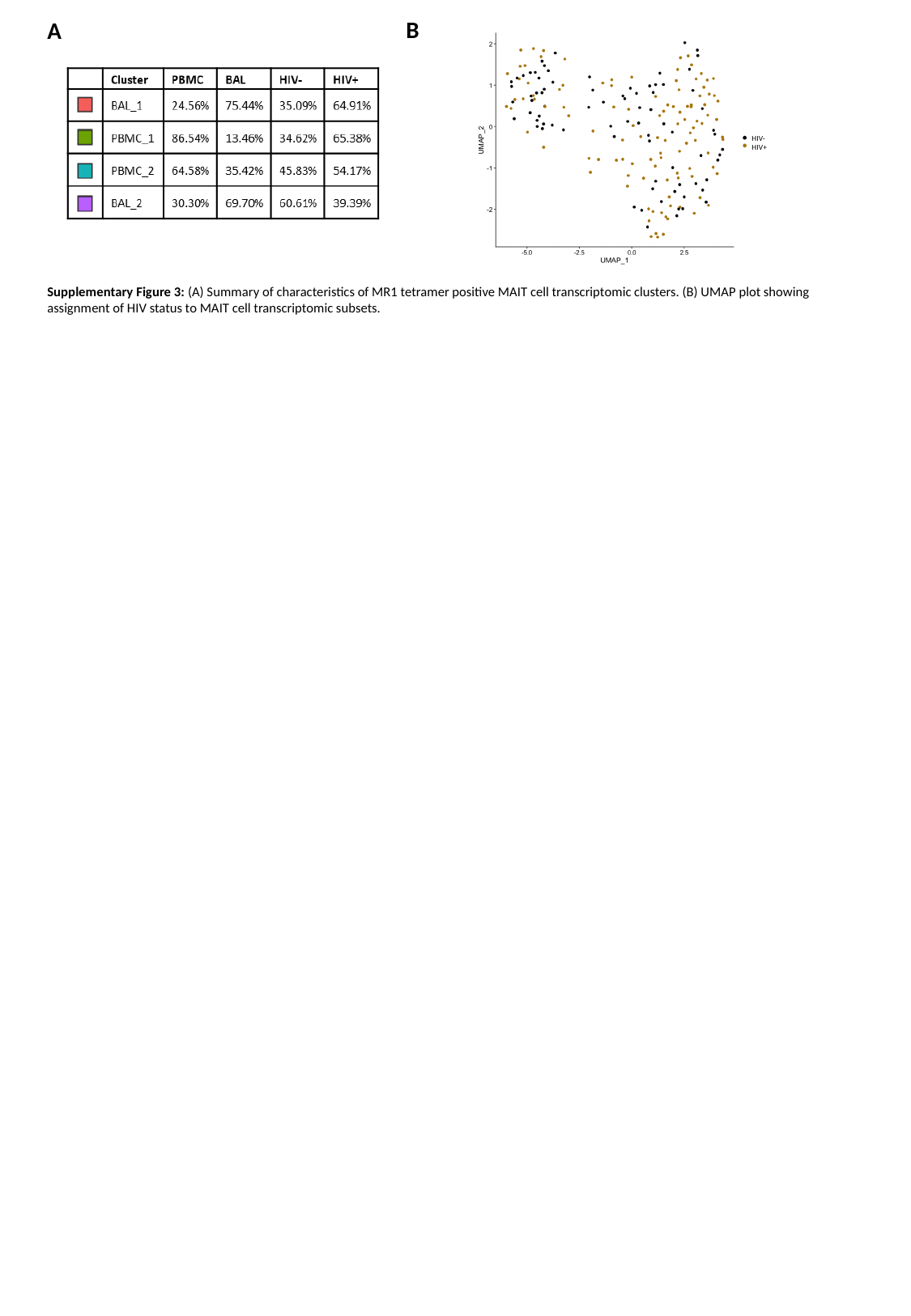

B
A
HIV-
HIV+
Supplementary Figure 3: (A) Summary of characteristics of MR1 tetramer positive MAIT cell transcriptomic clusters. (B) UMAP plot showing assignment of HIV status to MAIT cell transcriptomic subsets.

### Slide 4
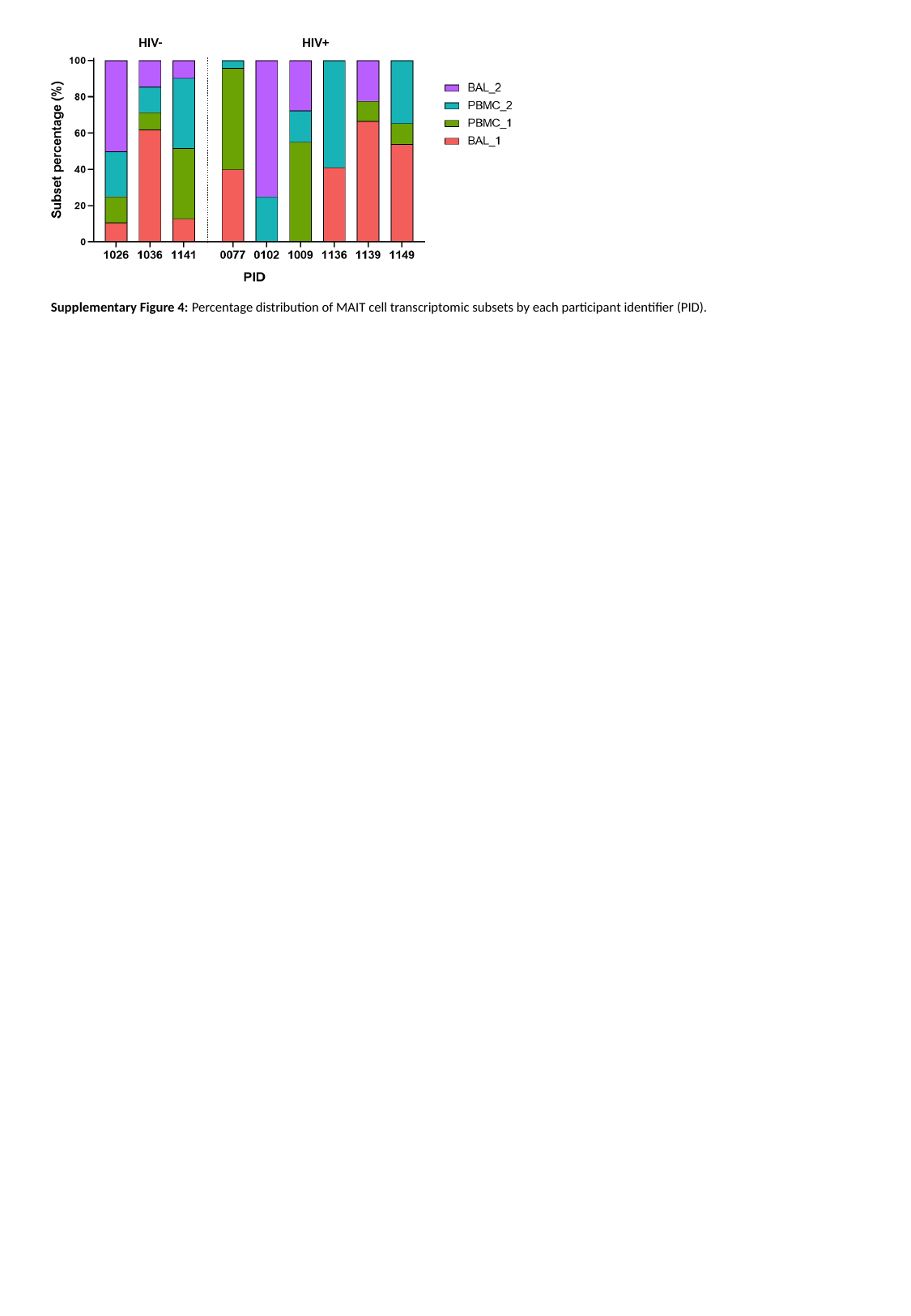

HIV-
HIV+
Supplementary Figure 4: Percentage distribution of MAIT cell transcriptomic subsets by each participant identifier (PID).

### Slide 5
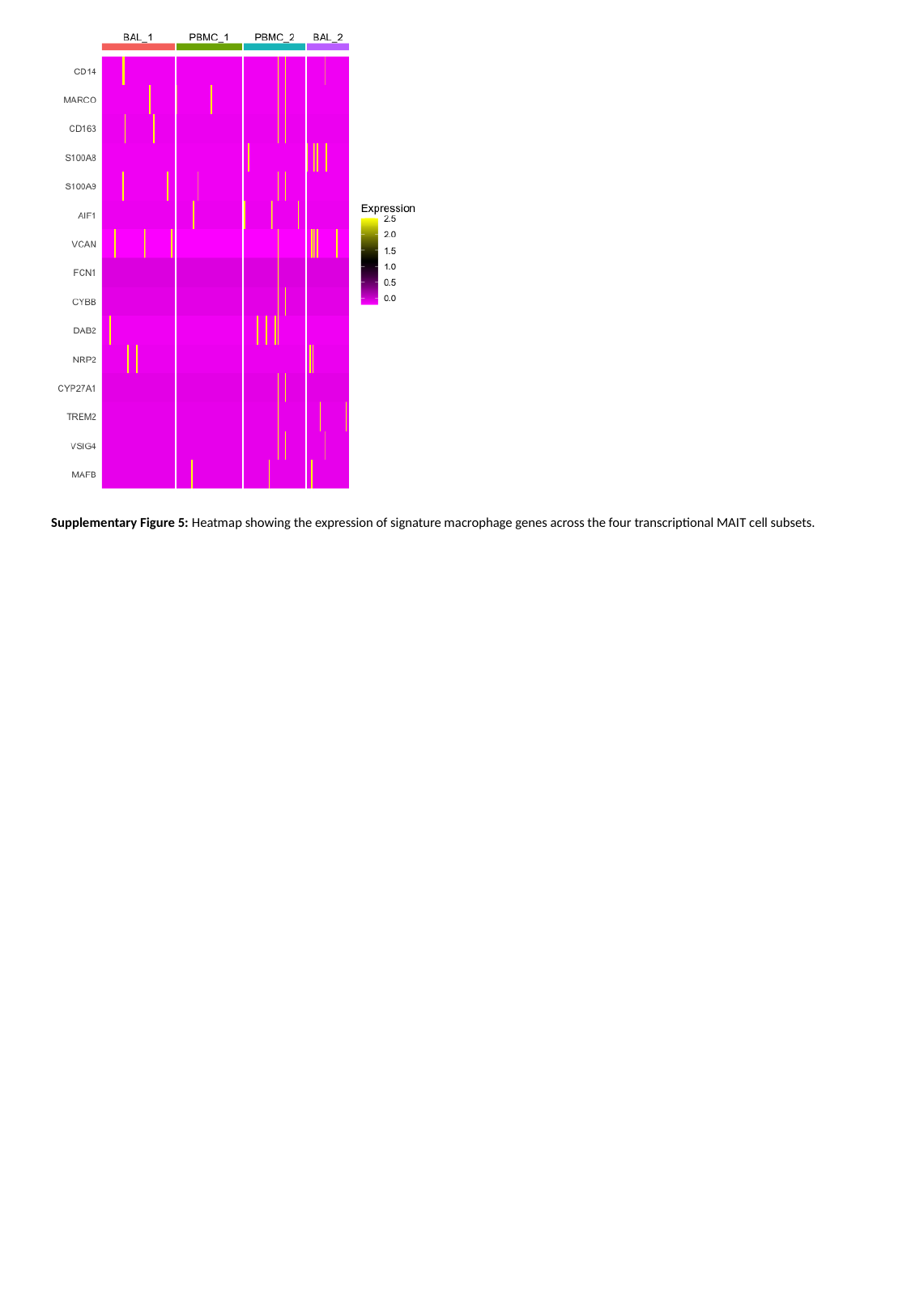

Supplementary Figure 5: Heatmap showing the expression of signature macrophage genes across the four transcriptional MAIT cell subsets.
